# Will The Grass Be Greener On The Other Side Of Climate Change?

**DOI:** 10.1101/2022.04.22.489016

**Authors:** Craig D Morris, Kevin P Kirkman, Pete JK Zacharias

## Abstract

Increasing atmospheric [CO_2_] is stimulating photosynthesis and plant production, increasing the demand for nitrogen relative to soil supply with declining global foliar nitrogen concentrations as a consequence. The effect of such oligotrophication on the forage quality of sweetveld, mixedveld, and sourveld grasslands in South Africa, which support livestock production and native ungulates, are unknown. Soil characteristics and the herbage quality of an abundant grass are described from baseline historical (mid 1980s) data collected across a sweet-mixed-sour grassland gradient in KwaZulu-Natal. Sourveld occurred on the most acidic, dystrophic soils and exhibited a pronounced decline in leaf N, digestibility, and other macronutrients during winter, in sharp contrast to sweetveld, on nutrient-rich soils, where forage quality varied little seasonally. In a carbon enriched, warmer, and most likely drier future climate, we predict that forage quality will be little altered in sweetveld where soil nutrients and temperature are not limiting but that sourveld could become ‘sourer’ because soil nutrients will be inadequate to match higher plant production promoted by elevated [CO_2_] and warmer and longer growing seasons. Reassessing historical data and seasonal and spatial monitoring of forage quality will enable past and future impacts of climate change on grassland forage quality to be assessed.

**Significance:**

- Grassland forage quality will likely decline with elevated [CO_2_] and warming, particularly in sourveld.
- Climate change could deepen and widen the sourveld winter forage bottleneck, necessitating greater supplementary feeding of livestock.

## Introduction

Grasslands, including the C_4_ dominated grasslands of South Africa face an uncertain future in a rapidly changing climate. The ongoing rise in the atmospheric concentration of anthropogenically-derived C0_2_ could increase carbon sequestration, biomass production, and the water use efficiency of grasses^1,2^, while also favouring woody species and alien invasive plants^3^. Reduced rainfall coupled with more frequent and severe droughts will further limit the production of herbage for livestock and the wild herbivores that grasslands support.^4,5^ Another, largely unrecognised, threat to livestock production posed by elevated [C0_2_] is an insidious decline in forage quality globally because of an increasing limitation of soil N supply to grasses growing faster over an extended growing season in a warmer, carbon-enriched atmosphere.^6^ Diminishing foliar [N] could cascade through ecosystems, slowing protein flow from plants to insect and mammalian herbivores.^7^ Even small decreases in the protein content and digestibility of forage would adversely affect animal health, reproduction, and weight gains.^7,8^

The potential impacts of elevated [C0_2_] (eCO2)and other climate change drivers on forage quality will occur across a well-recognised and agronomically important spatiotemporal gradient in South Africa, from sweetveld through mixedveld to sourveld. Livestock on sourveld require supplementary feeds and licks for up to six months^9^ because of the marked reduction in forage quality resulting from the translocation of foliar nutrients to roots at the end of the growing season^10^, whereas forage quality remains sufficiently high to maintain livestock production throughout the year in sweetveld areas; mixedveld displays intermediary seasonal quality changes^11^. Generally, with notable exceptions^11^, ‘sour’ grassland occurs in cool areas where high rainfall favours high primary production, but nutrient supply is limited on dystrophic soils, whereas in the hotter, usually lower-lying, sweetveld areas, typically on base-rich soils, low and erratic rainfall rather than nutrient availability restricts grass productivity^11,12^. Anthropogenic climate change could substantially alter the sweet-sour forage quality gradient because temperature, soil moisture and [C0_2_] interact to determine the balance between carbon assimilation and soil nutrient availability (particularly size of the mineralizable soil N pool) that determines spatial and seasonal differences in forage productivity and quality.^12,13^

To assess whether sourveld is becoming more ‘sour’ and sweetveld less ‘sweet’ owing to eC02 and other climate change drivers, we present historical plant quality data (collected in 1985-1986) to (1) describe soil physico-chemical and plant foliar nutrient gradients across sweet-mixed-sourveld sites, and (2) provide a baseline for detecting any oligotrophication that may have already occurred over the last third of a century. Atmospheric [C0_2_] has risen by more than 20% (346 to 418 ppm) since the mid-1980s (https://gml.noaa.gov/ccgg/trends/), during which spatially variable temperature increases in mean annual temperature^14^ of 0.01°C yr^−1^ to 0.03°C yr^−1^ and mean annual precipitation changes^15^ ranging from minus 12 mm to positive 14 mm have been recorded over South Africa. These trends are likely to accelerate because the southern African region is a global hotspot of climate change.^16^ We also consider uncertainties in the future likely trajectories of forage quality shifts in South African grasslands.

### Methods

Plant quality and soil characteristics were assessed at 31 sites across a sweet-sour grassland gradient in KwaZulu-Natal (Supplementary table 1).^17^ From 1985 to 1986, each site was visited at ca 73-day intervals to harvest foliar material of a consistent age - the top two leaves and a bud of vegetative tillers - from 20 plants of *Themeda triandra*, an abundant grass at all sites. We also refer to a wider study of winter (July) grass quality in the grassland biome in other provinces that used the same methods.^18^ Expert knowledge and literature were used to classify sites as ‘sweet’, ‘sour’ and ‘mixed’ types.

Plant quality analyses included for cellulase dry matter digestibility (%), leaf nitrogen concentration (%), chemical elemental analysis (N, P, K, Ca, Mg, Na, Zn, and S) (Supplementary table 2).

Topsoil was sampled (to 200 mm depth) from 20 combined auger points and assessed for particle size (texture), organic matter, field moisture capacity, pH, exchangeable acidity, acid saturation, effective cation exchange capacity, and P, K, Ca, Mg, Na (Supplementary table 3).

Forage quality (all seasons) and soil physico-chemical gradients in KwaZulu-Natal were examined using Principal Component Analysis of cross-correlation matrices, and seasonal differences in N% between grassland types were assessed with permutation ANOVAs (9999 permutations).

## Results

There was a strong forage quality gradient along which sourveld sites were most distinct for their low leaf digestibility, N%, and cation concentrations (Figure 1). Digestibility doubled and N% ranged five-fold across this gradient (Fig. 1 b & c).

**Figure 1:**
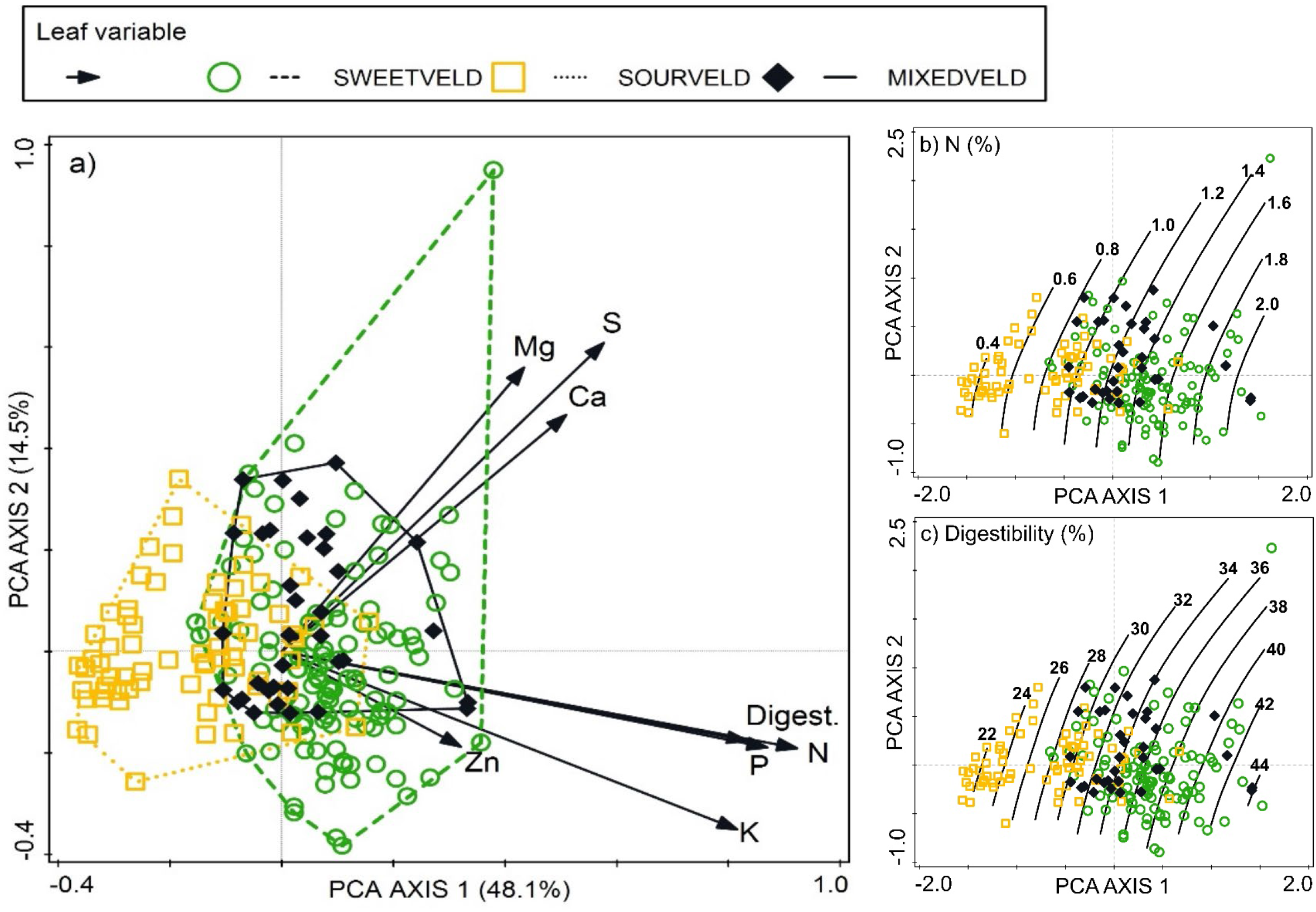
First two axes of a principal component analysis (PCA) of leaf variables measured at 31 locations in three grassland types in KwaZulu-Natal (a), and trend in leaf nitrogen (b) and digestibility (c) across the ordination.

Sweetveld soils were less acidic, had lower organic matter and capacity to hold water, but had substantially more exchangeable cations available than those from sourveld; mixedveld sites were intermediate (Figure 2).

**Figure 2:**
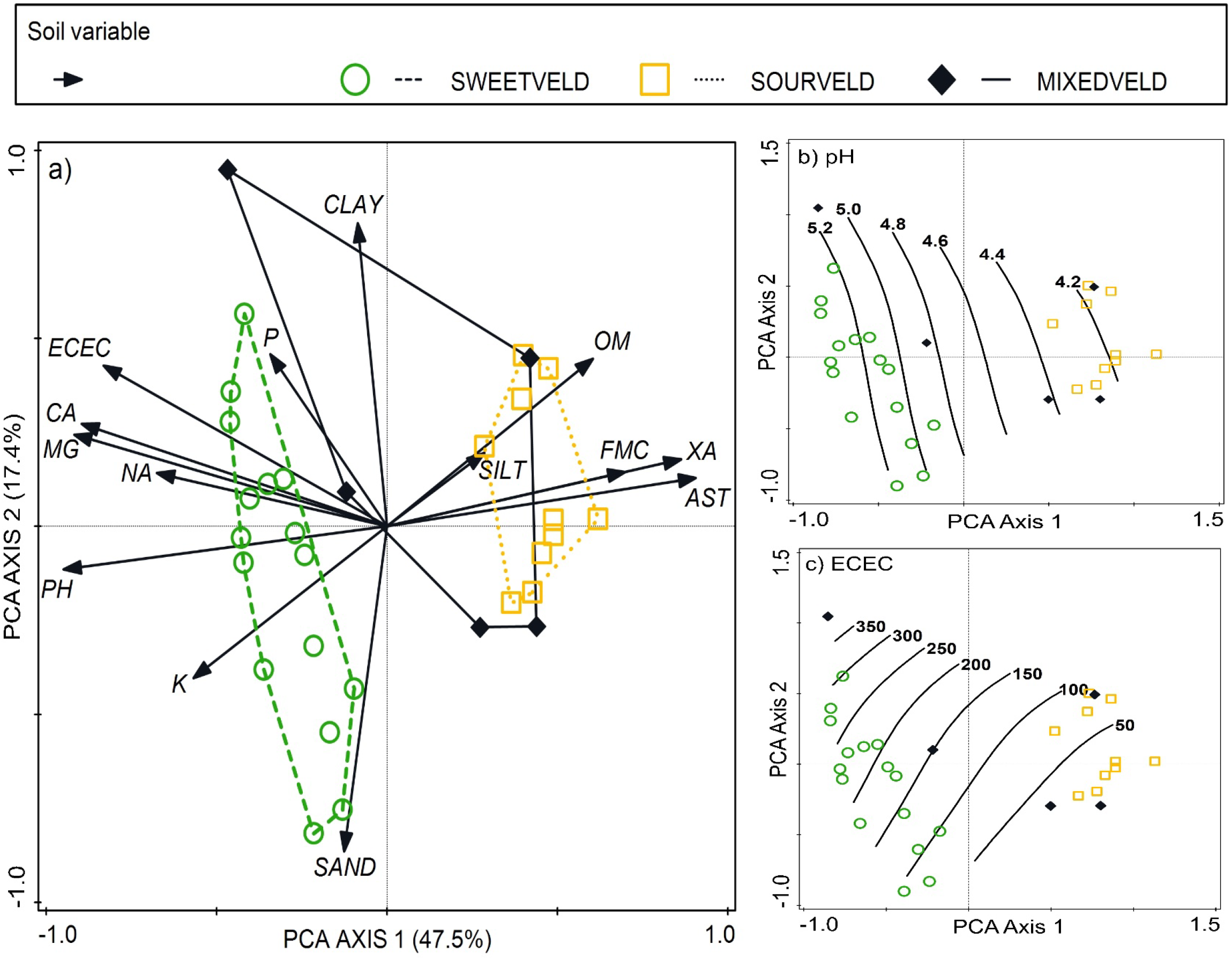
First two axes of a principal component analysis (PCA) of topsoil variables and trend in leaf measured at 31 locations in three grassland types in KwaZulu-Natal (a), and trend in pH (b) and ECEC (c) across the ordination. AST = acid saturation, ECEC = effective cation exchange capacity, FMC = field moisture capacity, OM = organic matter, XA =exchangeable acidity.

Sweetveld had consistently high foliar N%, declining only somewhat towards the winter of the second sampling season (Figure 3). In contrast, N concentration, similar to digestibility^17^, declined to markedly low levels from late summer through to early spring. Quality trends for mixedveld were not consistent nor pronounced. Summer and winter levels of N above 1.0% and below 0.5%, respectively for sweetveld and sourveld in KwaZulu-Natal matched the winter extremes measured elsewhere in the grassland biome^18^. Also seasonally variable in sourveld and to a lesser extent in mixedveld were leaf concentrations of the macronutrients, P, K, Mg, and S.^17^

**Figure 3:**
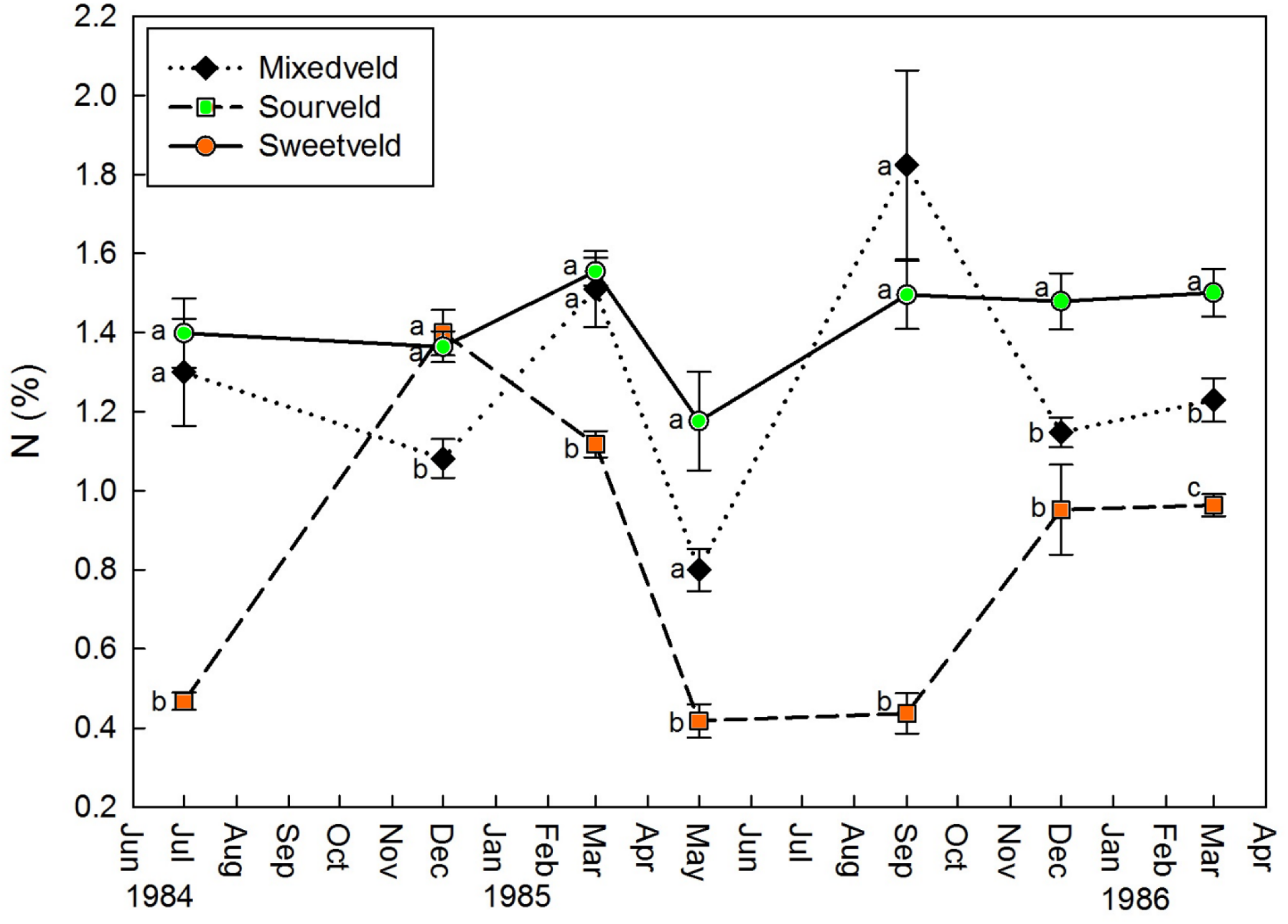
Seasonal trends in mean (±se) leaf nitrogen concentration (%) measured in three grassland types in KwaZulu-Natal. Means within sampling dates with letters in common are not different (p = 0.05).

The links between forage quality, soil characteristics and environment were weak, with increasing altitude the only consistent predictor of ‘sourness’ in KwaZulu-Natal.^17,18^

## Discussion

Future reductions in the forage quality of sourveld and sweetveld will depend on how other major climate change drivers (temperature and precipitation) interact with the carbon fertilisation effect (CFE) to alter the balance between plant growth-driven demand for, and soil supply of, nutrients, primarily N.^6,7,12^ The CFE would be most pronounced when resources and environmental conditions do not restrict plant growth.^19^ However, the effects of multi-way interactions between climate drivers on plants and soils, particular on forage quality, are poorly understood; these interactions can be complex, multiplicative^20,21^, and species-specific^22^. Given these uncertainties and the current growth limitations prevailing in sour- and sweetveld^11,12^, we tentatively predict the following potential shifts in plant quality (leaf [N], digestibility, and fibre content) under climate change.

Sweetveld mostly occurs in semi-arid regions, where soil moisture, not nutrients, limit plant growth.^12^ Despite lower rainfall and increased evaporative demand with warming predicted for semi-arid regions^5^, plant growth could increase because of more efficient water use under eCO2 resulting from stomal closure^1,22^. Elevated temperatures combined with the CFE could stimulate grass production, lowering the N content and digestibility of herbage.^8^ However, extreme and prolonged droughts and heat waves, both of which will become more frequent with climate change^5,16^, will limit carbon assimilation and nutrient availability by curtailing microbial decomposition and nutrient cycling^19,20^. The consequences of climate change in semi-arid regions are still uncertain^21^ but it is not likely that sweetveld will experience a consistent large directional change in productivity and forage quality in the future.

In sourveld, predominately in higher lying, cooler, moister climes, nutrient-poor soils and low temperatures act together to constrain the quality of forage during the non-growing season^11,12^. Warming, especially in spring and autumn, and eCO_2_ would likely enable a longer period of growth, further limiting soil nutrient supply^6^ and reducing winter forage quality. Increased N mineralisation with warming could mitigate this potential future decline in forage quality on organic mountain soils^23^ but not on the dystrophic mineral soils of outlier sourveld areas at low elevations^11^.

The critical winter forage bottleneck in forage quality in sourveld (Figure 3) is likely to be exacerbated in the future because eCO_2_-driven reductions in protein content and digestibility in late summer and autumn occur when nutrient in senescing plants are already below critical levels for livestock^9,24^. Consequently, supplementary feeding costs will increase^8^ while wild ungulates would need to forage differently to match their metabolic requirements^25^. Higher-quality C_3_ grasses that remain greener for longer in autumn could obtain a competitive advantage over C_4_ species in the future^26^ but only at the higher and far western margins of the grassland biome, and perhaps not to any significant extent^2^.

Research is required across a sweet-mixed-to-sourveld gradient to understand patterns and mechanisms of seasonal nutrient flows between plant parts – our knowledge of these is still surprisingly rudimentary^10^, – species-specific responses to interactions between multiple climate change drivers^22^, the potential effects on plant growth and quality of ongoing atmospheric nitrogen deposition^20^, and the extent to which the CFE effect on plant quality could be attenuated by downregulation of photosynthesis through acclimation and progressive N limitation over time^1,7,8^. We also recommend resampling sites with historical plant data, such as those presented here (Supplementary tables 1 & 2), to establish the extent to which CFE-driven shifts in forage quality may have already occurred over the last few decades, and to monitor regularly, widely, and seasonally shifts in leaf stoichiometry (at minimum C : N ratios^7,13^) to establish the degree and extent of climate-driven oligotrophication^8^.

## Conclusion

Climate change has the potential to alter in agronomically important ways the current spatial and seasonal patterns of grass forage quality in South African grasslands. We predict the greatest ‘souring’ will occurring in sourveld, with a minimal response in sweetveld, but there are many uncertainties as to the direction and rate of change in forage quality and the extent to which such changes will affect livestock production and wild ungulates. Further detailed research and regular monitoring are required to assess if, where, how, and why forage quality of grasslands in South Africa is responding to climate change.

## REFERENCES

1. Lee M, Manning P, Rist J, Power SA, Marsh C. A global comparison of grassland biomass responses to CO2 and nitrogen enrichment. Philos. Trans. R. Soc. Lond., B, Biol. Sci. 2010;365(1549):2047–2056. https://doi.org/10.1098/rstb.2010.0028

2. Wand SJ, Midgley GF, Jones MH, Curtis PS. Responses of wild C4 and C3 grass (Poaceae) species to elevated atmospheric CO2 concentration: a meta-analytic test of current theories and perceptions. Glob. Change Biol. 1999;5(6):723–741. https://doi.org/10.2989/10220119.2014.939996

3. O’connor TG, Puttick JR, Hoffman MT. Bush encroachment in southern Africa: changes and causes. Afr. J. Range Forage Sci. 2014;31(2):67–88. https://doi.org/10.1007/978-81-322-2265-1_7

4. Giridhar K, Samireddypalle A. Impact of climate change on forage availability for livestock. In: Sejian V, Gaughan J, Baumgard L, Prasad C, editors. Climate change impact on livestock: adaptation and mitigation. New Delhi: Springer; 2015. pp. 97–112.

5. Davis CL, Vincent K. Climate risk and vulnerability: a handbook for Southern Africa, 2nd ed. 2017. Pretoria: CSIR, South Africa; 2017. Available at: https://researchspace.csir.co.za/dspace/handle/10204/10066

6. Craine JM, Elmore AJ, Wang L, Aranibar J, Bauters M, Boeckx P et al. Isotopic evidence for oligotrophication of terrestrial ecosystems. Nat. Ecol. Evol. 2018;2(11):1735–1744. https://doi.org/10.1038/s41559-018-0694-0

7. Mason RE, Craine JM, Lany NK, Jonard M, Ollinger SV, Groffman PM et al. Evidence, causes, and consequences of declining nitrogen availability in terrestrial ecosystems. Science 2022;376(6590):eabh3767. https://doi.org/10.1126/science.abh3767

8. Augustine DJ, Blumenthal DM, Springer TL, LeCain DR, Gunter SA, Derner JD. Elevated CO2 induces substantial and persistent declines in forage quality irrespective of warming in mixedgrass prairie. Ecol. Appl 2018;28(3):721–735. https://doi.org/10.1002/eap.1680

9. Kirkman KP, Moore A. Perspective: towards improved grazing management recommendations for sourveld. Afr. J. Range Forage Sci. 12(3):135–44. https://doi.org/10.1080/10220119.1995.9647883

10. Weinmann H. Seasonal chemical changes in the roots of some South African highveld grasses. J. S. African Bot. 1940:131–145.

11. Tainton NM. The production characteristics of the main grazing lands of South Africa. In: Tainton NM, editor. Veld Management in South Africa. Pietermaritzburg: University of Natal Press; 1999. p. 46–52.

12. Ellery WN, Scholes RJ, Scholes MC. The distribution of sweetveld and sourveld in South Africa’s grassland biome in relation to environmental factors. Afr. J. Range Forage Sci. 1995;12(1):38–45. https://doi.org/10.1080/10220119.1995.9647860

13. Kunz RP, Schulze RE, Scholes RJ. An approach to modelling spatial changes of plant carbon: nitrogen ratios in southern Africa in relation to anticipated global climate change. J. Biogeogr. 1995;1(2/3):401–408. https://doi.org/10.2307/2845936

14. Jury MR. Climate trends across South Africa since 1980. Water S.A. 2018;44(2):297–307. https://doi.org/10.4314/wsa.v44i2.15

15. Marumbwa FM, Cho MA, Chirwa PW. Analysis of spatio-temporal rainfall trends across southern African biomes between 1981 and 2016. Phys. Chem. Earth A/B/C. 2019;114(Dec):102808. https://doi.org/10.1016/j.pce.2019.10.004

16. Engelbrecht FA, Monteiro P. The IPCC Assessment Report Six Working Group 1 report and southern Africa: Reasons to take action. S. Afr. J. Sci. 2021;117(11-12):1–7. https://doi.org/10.17159/sajs.2021/12679

17. Zacharias PJ. The seasonal patterns in plant quality in various ecological zones in Natal [MSc thesis]. Pietermaritzburg: University of Natal; 1990. Available at: https://researchspace.ukzn.ac.za/handle/10413/20233

18. Kirkman KP. Factors affecting the seasonal variation of veld quality in South Africa [MSc thesis]. Pietermaritzburg: University of Natal; 1988. Available at: https://researchspace.ukzn.ac.za/handle/10413/20232

19. Seibert R, Donath TW, Moser G, Laser H, Grünhage L, Schmid T et al. Effects of long-term CO2 enrichment on forage quality of extensively managed temperate grassland. Agric. Ecosyst. Environ. 2021;312:107347. https://doi.org/10.1016/j.agee.2021.107347

20. Polley HW, Morgan JA, Fay PA. Application of a conceptual framework to interpret variability in rangeland responses to atmospheric CO2 enrichment. J. Agric. Sci. 2011;149(1):1–4. https://doi.org/10.1017/s0021859610000717

21. Song J, Wan S, Piao S, Knapp AK, Classen AT, Vicca S et al. A meta-analysis of 1,119 manipulative experiments on terrestrial carbon-cycling responses to global change. Nat. Ecol. Evol. 2019;3(9):1309–1320. https://doi.org/10.1038/s41559-019-0958-3

22. Pastore MA, Lee TD, Hobbie SE, Reich PB. Interactive effects of elevated CO2, warming, reduced rainfall, and nitrogen on leaf gas exchange in five perennial grassland species. Plant Cell Environ. 2020;43(8):1862–1878. https://doi.org/10.1111/pce.13783

23. Carbutt C, Edwards TJ, Fynn RW, Beckett RP. Evidence for temperature limitation of nitrogen mineralisation in the Drakensberg Alpine Centre. S. Afr. J. Bot. 2013;88:447–54. https://doi.org/10.1016/j.sajb.2013.09.001

24. Milchunas DG, Mosier AR, Morgan JA, LeCain DR, King JY, Nelson JA. Elevated CO2 and defoliation effects on a shortgrass steppe: forage quality versus quantity for ruminants. Agric. Ecosyst. Environ. 2005;111(1-4):166–84. https://doi.org/10.1016/j.agee.2005.06.014

25. Searle KR, Hobbs NT, Gordon IJ. It’s the “foodscape”, not the landscape: using foraging behavior to make functional assessments of landscape condition. Isr. J. Ecol. Evol. 2007;53(3-4):297–316. https://doi.org/10.1560/ijee.53.3.297

26. Chamaillé-Jammes S, Bond WJ. Will global change improve grazing quality of grasslands? A call for a deeper understanding of the effects of shifts from C4 to C3 grasses for large herbivores. Oikos 2010;119(12):1857–61. https://doi.org/10.1111/j.1600-0706.2010.19070.x

