## Supplementary tables 1-3 for "Will The Grass Be Greener On The Other Side Of Climate Change?"

**Table 1:** Location and physiography of sample sites (data from Zacharias 1990)

| Site # | Location | Latitude (S) | Longitude (E) | Altitude (m asl) | Aspect (°) | Slope (°) | Classification |
|---|---|---|---|---|---|---|---|
| 2 | Umfolozi, HiP[1] | 28°14'20" | 31°48'20" | 145 | 26 | 5 | Sweetveld |
| 3 | Umfolozi, HiP | 28°19'00" | 31°50'20" | 274 | 68 | 9 | Sweetveld |
| 4 | Umfolozi, HiP | 28°19'10" | 31°50'20" | 259 | 26 | 8 | Sweetveld |
| 5 | Umfolozi, HiP | 28°19'20" | 31°50'20" | 332 | 0 | 0 | Sweetveld |
| 6 | Hluhluwe, HiP | 28°06'10" | 32°02'30" | 305 | 74 | 0 | Sweetveld |
| 7 | Hluhluwe, HiP | 28°05'06" | 32°07'25" | 122 | 180 | 5 | Sweetveld |
| 8 | Mgambo Farm, Magudu | 27°30'35" | 31°40'10" | 419 | 85 | 2 | Sweetveld |
| 9 | Mgambo Farm, Magudu | 27°30'35" | 31°40'12" | 305 | 95 | 6 | Sweetveld |
| 10 | Mgambo Farm, Magudu | 27°30'50" | 31°40'00" | 427 | 115 | 7 | Sweetveld |
| 11* | Ronberbosch Farm, Magudu | 27°46'40" | 31°39'10" | 488 | 130 | 5 | Sweetveld |
| 12 | Mkuze Game Reserve | 27°38'40" | 32°09'20" | 122 | 140 | 15 | Sweetveld |
| 13 | Mkuze Game Reserve | 27°38'40" | 32°09'30" | 183 | 40 | 12 | Sweetveld |
| 14 | Mkuze Game Reserve | 27°38'30" | 32°09'45" | 183 | 0 | 0 | Sweetveld |
| 15 | Mkuze Game Reserve | 27°36'40" | 32°13'15" | 122 | 0 | 0 | Sweetveld |
| 16 | Mkuze Game Reserve | 27°39'50" | 32°14'30" | 61 | 85 | 4 | Sweetveld |
| 17 | Mkuze Game Reserve | 27°40'00" | 32°15'00" | 61 | 0 | 0 | Sweetveld |
| 18 | Hiighmoor Nature Reserve, UDP[2] | 29°19'15" | 29°37'25" | 2024 | 4 | 12 | Sourveld |
| 19* | Hiighmoor Nature Reserve, UDP | 29°15'00" | 29°38'40" | 1844 | 0 | 15 | Sourveld |
| 20 | Hiighmoor Nature Reserve, UDP | 29°18'55" | 29°38'50" | 1814 | 145 | 19 | Sourveld |
| 21 | Hiighmoor Nature Reserve, UDP | 29°19'30" | 29°37'10" | 2057 | 60 | 12 | Sourveld |
| 22 | Cathedral Peak, UDP | 28°57'40" | 29°14'20" | 1768 | 175 | 4 | Sourveld |
| 23 | Cathedral Peak, UDP | 28°57'40" | 29°14'05" | 1783 | 35 | 15 | Sourveld |
| 24 | Cathedral Peak, UDP | 28°58'20" | 29°15'05" | 1844 | 0 | 0 | Sourveld |
| 25 | Cathedral Peak, UDP | 28°58'55" | 29°14'30" | 1899 | 151 | 15 | Sourveld |
| 26 | Cathedral Peak, UDP | 28°58'55" | 29°14'31" | 1899 | 10 | 15 | Sourveld |
| 27 | Cathedral Peak, UDP | 28°58'00" | 29°16'00" | 2057 | 60 | 28 | Sourveld |
| 28 | Ukulinga Research Farm, Pietermaritzburg | 29°39'48" | 30°24'08" | 820 | 90 | 10 | Mixedveld |
| 29 | Ukulinga Research Farm, Pietermaritzburg | 29°40'06" | 30°24'15" | 840 | 0 | 0 | Mixedveld |
| 30 | Worlds View, Pietermaritzburg | 29°34'49" | 30°20'00" | 884 | 80 | 15 | Mixedveld |
| 31* | Kranskloof Nature Reserve, Kloof | 29°45'18" | 30°50'06" | 152 | 180 | 26 | Mixedveld |
| 32 | Kranskloof Nature Reserve, Kloof | 29°45'52" | 30°51'23" | 122 | 160 | 13 | Mixedveld |

*Location uncertain.

[1]HiP = Hluhluwe–iMfolozi Park

[2]UDP = uKhahlamba-Drakensberg Park

Note: Site 1 was destroyed by Tropical Storm Domoina in 1984.

**Supplementary Table 1:** Digestibility and nutrient content of *Themeda triandra* leaves sampled at 31 sites from July 1984 to March 1986 in KwaZulu-Natal (data from Zacharias 1990)

| Site # | LOCATION | Month | Year | Digestibility (%) | N (%) | P (%) | K (%) | Ca (%) | Mg (%) | S (%) | Zn (mg/kg) |
|---|---|---|---|---|---|---|---|---|---|---|---|
| 2 | UMFOLOZI | Jul | 1984 | 29.31 | 0.99 | 0.06 | 0.61 | 0.15 | 0.3 | 0.13 | 25 |
| 2 | UMFOLOZI | Dec | 1984 | 33.58 | 1.03 | 0.08 | 1.01 | 0.16 | 0.39 | 0.13 | 17 |
| 2 | UMFOLOZI | Mar | 1985 | 34.96 | 1.51 | 0.11 | 1.25 | 0.16 | 0.35 | 0.15 | 21 |
| 2 | UMFOLOZI | May | 1985 | 29.1 | 0.18 | 0.04 | 0.75 | 0.19 | 0.31 | 0.14 | 25 |
| 2 | UMFOLOZI | Sep | 1985 | 36.7 | 0.73 | 0.06 | 0.97 | 0.2 | 0.31 | 0.19 | 18 |
| 2 | UMFOLOZI | Dec | 1985 | 42.08 | 1.31 | 0.09 | 1.17 | 0.21 | 0.33 | 0.23 | 16 |
| 2 | UMFOLOZI | Mar | 1986 | 41.38 | 2.01 | 0.15 | 1.14 | 0.25 | 0.36 | 0.23 | 25 |
| 3 | UMFOLOZI | Jul | 1984 | 28.9 | 0.97 | 0.06 | 0.51 | 0.17 | 0.3 | 0.15 | 20 |
| 3 | UMFOLOZI | Dec | 1984 | 38.8 | 1.39 | 0.11 | 1.14 | 0.21 | 0.34 | 0.13 | 14 |
| 3 | UMFOLOZI | Mar | 1985 | 36.42 | 1.56 | 0.11 | 1.17 | 0.17 | 0.38 | 0.13 | 19 |
| 3 | UMFOLOZI | May | 1985 | 28.14 | 0.95 | 0.03 | 0.64 | 0.31 | 0.42 | 0.17 | 17 |
| 3 | UMFOLOZI | Sep | 1985 | 46.96 | 1.85 | 0.08 | 1.83 | 0.24 | 0.44 | 0.16 | 15 |
| 3 | UMFOLOZI | Dec | 1985 | 36.45 | 1.19 | 0.07 | 1.3 | 0.2 | 0.3 | 0.11 | 13 |
| 3 | UMFOLOZI | Mar | 1986 | 40.67 | 1.5 | 0.09 | 1.25 | 0.31 | 0.3 | 0.14 | 21 |
| 4 | UMFOLOZI | Jul | 1984 | 29.2 | 1.04 | 0.06 | 0.6 | 0.3 | 0.4 | 0.13 | 20 |
| 4 | UMFOLOZI | Dec | 1984 | 35.95 | 1.42 | 0.09 | 1.02 | 0.23 | 0.3 | 0.12 | 20 |
| 4 | UMFOLOZI | Mar | 1985 | 35.21 | 1.59 | 0.11 | 1.21 | 0.22 | 0.33 | 0.13 | 20 |
| 4 | UMFOLOZI | May | 1985 | 32.44 | 0.71 | 0.04 | 0.83 | 0.19 | 0.47 | 0.18 | 15 |
| 4 | UMFOLOZI | Sep | 1985 | 49.88 | 1.59 | 0.1 | 1.8 | 0.24 | 0.51 | 0.2 | 14 |
| 4 | UMFOLOZI | Dec | 1985 | 36.87 | 1.24 | 0.09 | 1.47 | 0.2 | 0.35 | 0.16 | 15 |
| 4 | UMFOLOZI | Mar | 1986 | 41.79 | 1.5 | 0.13 | 1.25 | 0.22 | 0.28 | 0.12 | 23 |
| 5 | UMFOLOZI | Jul | 1984 | 26.41 | 1.11 | 0.07 | 0.5 | 0.2 | 0.43 | 0.15 | 18 |
| 5 | UMFOLOZI | Dec | 1984 | 40.44 | 1.38 | 0.12 | 1.14 | 0.24 | 0.3 | 0.14 | 17 |
| 5 | UMFOLOZI | Mar | 1985 | 32.51 | 1.64 | 0.11 | 1.25 | 0.19 | 0.31 | 0.14 | 20 |
| 5 | UMFOLOZI | May | 1985 | 33.8 | 0.82 | 0.06 | 0.91 | 0.25 | 0.42 | 0.18 | 13 |
| 5 | UMFOLOZI | Sep | 1985 | 53.62 | 0.97 | 0.07 | 1.1 | 0.21 | 0.36 | 0.17 | 13 |
| 5 | UMFOLOZI | Dec | 1985 | 36.9 | 1.2 | 0.09 | 1.28 | 0.19 | 0.32 | 0.16 | 15 |
| 5 | UMFOLOZI | Mar | 1986 | 40.04 | 1.53 | 0.1 | 1.37 | 0.24 | 0.36 | 0.14 | 21 |
| 6 | HLUHLUWE | Jul | 1984 | 26.83 | 1.31 | 0.08 | 0.5 | 0.27 | 0.55 | 0.17 | 20 |
| 6 | HLUHLUWE | Dec | 1984 | 34.79 | 0.99 | 0.1 | 0.99 | 0.14 | 0.26 | 0.1 | 16 |
| 6 | HLUHLUWE | Mar | 1985 | 34.81 | 1.36 | 0.11 | 1.05 | 0.16 | 0.32 | 0.12 | 18 |
| 6 | HLUHLUWE | May | 1985 | 29.26 | 1 | 0.07 | 0.87 | 0.31 | 0.41 | 0.15 | 23 |
| 6 | HLUHLUWE | Sep | 1985 | 40.89 | 1.57 | 0.13 | 1.77 | 0.25 | 0.36 | 0.15 | 20 |
| 6 | HLUHLUWE | Dec | 1985 | 33.58 | 1.14 | 0.1 | 1.04 | 0.17 | 0.24 | 0.13 | 15 |
| 6 | HLUHLUWE | Mar | 1986 | 37.34 | 1.29 | 0.1 | 1.01 | 0.2 | 0.38 | 0.12 | 15 |
| 7 | HLUHLUWE | Jul | 1984 | 30.74 | 1.04 | 0.18 | 0.64 | 0.26 | 0.56 | 0.14 | 22 |
| 7 | HLUHLUWE | Dec | 1984 | 35.68 | 1.55 | 0.16 | 1.17 | 0.14 | 0.31 | 0.11 | 19 |
| 7 | HLUHLUWE | Mar | 1985 | 33.89 | 1.38 | 0.15 | 1.1 | 0.15 | 0.25 | 0.09 | 21 |
| 7 | HLUHLUWE | May | 1985 | 26.91 | 1.18 | 0.18 | 0.92 | 0.26 | 0.41 | 0.12 | 18 |
| 7 | HLUHLUWE | Sep | 1985 | 38.55 | 1.82 | 0.2 | 1.64 | 0.24 | 0.29 | 0.12 | 22 |
| 7 | HLUHLUWE | Dec | 1985 | 33.1 | 1.3 | 0.16 | 1.16 | 0.2 | 0.22 | 0.09 | 19 |
| 7 | HLUHLUWE | Mar | 1986 | 35.29 | 1.15 | 0.14 | 0.9 | 0.17 | 0.34 | 0.09 | 16 |
| 8 | MGAMBO FARM | Jul | 1984 | 38.41 | 1.85 | 0.2 | 0.92 | 0.25 | 0.45 | 0.15 | 24 |
| 8 | MGAMBO FARM | Dec | 1984 | 37.68 | 1.55 | 0.19 | 1.38 | 0.18 | 0.34 | 0.14 | 18 |
| 8 | MGAMBO FARM | Mar | 1985 | 38.41 | 1.53 | 0.14 | 1.17 | 0.18 | 0.26 | 0.11 | 20 |
| 8 | MGAMBO FARM | May | 1985 | 29.93 | 1.08 | 0.09 | 1.08 | 0.2 | 0.32 | 0.14 | 17 |
| 8 | MGAMBO FARM | Sep | 1985 | 35.8 | 1.4 | 0.11 | 1.16 | 0.2 | 0.35 | 0.14 | 17 |
| 8 | MGAMBO FARM | Dec | 1985 | 41.84 | 1.71 | 0.14 | 1.29 | 0.21 | 0.36 | 0.14 | 18 |

| | | | | | | | | | | | |
|---|---|---|---|---|---|---|---|---|---|---|---|
| 8 | MGAMBO FARM | Mar | 1986 | 40.33 | 1.62 | 0.15 | 1.23 | 0.19 | 0.27 | 0.12 | 21 |
| 9 | MGAMBO FARM | Jul | 1984 | 38.27 | 1.91 | 0.17 | 1.01 | 0.21 | 0.46 | 0.15 | 22 |
| 9 | MGAMBO FARM | Dec | 1984 | 37.65 | 1.39 | 0.11 | 1.36 | 0.14 | 0.32 | 0.12 | 19 |
| 9 | MGAMBO FARM | Mar | 1985 | 34.9 | 1.62 | 0.14 | 1.22 | 0.17 | 0.3 | 0.13 | 22 |
| 9 | MGAMBO FARM | May | 1985 | 25.73 | 0.73 | 0.06 | 0.7 | 0.16 | 0.29 | 0.14 | 18 |
| 9 | MGAMBO FARM | Sep | 1985 | 35.5 | 1.43 | 0.11 | 1.13 | 0.19 | 0.34 | 0.16 | 20 |
| 9 | MGAMBO FARM | Dec | 1985 | 44.1 | 2.1 | 0.17 | 1.54 | 0.22 | 0.39 | 0.18 | 23 |
| 9 | MGAMBO FARM | Mar | 1986 | 36.64 | 1.7 | 0.15 | 1.28 | 0.18 | 0.13 | 0.14 | 23 |
| 10 | MGAMBO FARM | Jul | 1984 | 36.04 | 1.64 | 0.16 | 0.75 | 0.23 | 0.65 | 0.13 | 22 |
| 10 | MGAMBO FARM | Dec | 1984 | 40.82 | 1.31 | 0.11 | 0.97 | 0.18 | 0.31 | 0.11 | 20 |
| 10 | MGAMBO FARM | Mar | 1985 | 34.78 | 1.37 | 0.12 | 1.03 | 0.16 | 0.32 | 0.12 | 21 |
| 10 | MGAMBO FARM | May | 1985 | 31.51 | 1.71 | 0.15 | 1.58 | 0.25 | 0.5 | 0.13 | 20 |
| 10 | MGAMBO FARM | Sep | 1985 | 37.38 | 1.45 | 0.14 | 1.3 | 0.22 | 0.42 | 0.12 | 20 |
| 10 | MGAMBO FARM | Dec | 1985 | 43.26 | 1.2 | 0.12 | 1.02 | 0.19 | 0.33 | 0.12 | 21 |
| 10 | MGAMBO FARM | Mar | 1986 | 36.52 | 1.44 | 0.12 | 1.08 | 0.17 | 0.37 | 0.13 | 22 |
| 11 | RONDERBOSCH FARM | Jul | 1984 | 31.05 | 0.92 | 0.08 | 0.6 | 0.12 | 0.3 | 0.17 | 20 |
| 11 | RONDERBOSCH FARM | Dec | 1984 | 31.76 | 1.44 | 0.1 | 1.07 | 0.12 | 0.24 | 0.13 | 17 |
| 11 | RONDERBOSCH FARM | Mar | 1985 | 35.54 | 1.66 | 0.13 | 1.15 | 0.26 | 0.23 | 0.14 | 18 |
| 11 | RONDERBOSCH FARM | May | 1985 | 27.19 | 1.74 | 0.14 | 1.67 | 0.23 | 0.36 | 0.2 | 15 |
| 11 | RONDERBOSCH FARM | Sep | 1985 | 35.6 | 1.68 | 0.12 | 1.69 | 0.19 | 0.33 | 0.21 | 14 |
| 11 | RONDERBOSCH FARM | Dec | 1985 | 43.21 | 1.64 | 0.12 | 1.61 | 0.17 | 0.29 | 0.21 | 15 |
| 11 | RONDERBOSCH FARM | Mar | 1986 | 37.31 | 1.74 | 0.14 | 1.2 | 0.27 | 0.24 | 0.15 | 19 |
| 12 | MKUZE | Jul | 1984 | 35.12 | 1.42 | 0.11 | 0.84 | 0.31 | 0.4 | 0.22 | 18 |
| 12 | MKUZE | Dec | 1984 | 35.33 | 1.47 | 0.09 | 1.01 | 0.22 | 0.37 | 0.16 | 17 |
| 12 | MKUZE | Mar | 1985 | 35.99 | 1.65 | 0.1 | 1.13 | 0.35 | 0.28 | 0.14 | 19 |
| 12 | MKUZE | May | 1985 | 31.46 | 1.98 | 0.15 | 1.84 | 0.32 | 0.5 | 0.22 | 17 |
| 12 | MKUZE | Sep | 1985 | 37.23 | 1.13 | 0.06 | 1.15 | 0.25 | 0.34 | 0.14 | 19 |
| 12 | MKUZE | Dec | 1985 | 37.49 | 1.65 | 0.1 | 1.51 | 0.2 | 0.29 | 0.16 | 18 |
| 12 | MKUZE | Mar | 1986 | 37.85 | 1.48 | 0.11 | 1.26 | 0.23 | 0.35 | 0.19 | 21 |
| 13 | MKUZE | Jul | 1984 | 34.93 | 1.64 | 0.13 | 0.95 | 0.34 | 0.48 | 0.18 | 25 |
| 13 | MKUZE | Dec | 1984 | 36.71 | 1.35 | 0.12 | 1.01 | 0.2 | 0.45 | 0.13 | 16 |
| 13 | MKUZE | Mar | 1985 | 35.75 | 1.5 | 0.11 | 1.2 | 0.33 | 0.29 | 0.12 | 20 |
| 13 | MKUZE | May | 1985 | 28.93 | 1.95 | 0.17 | 1.95 | 0.35 | 0.45 | 0.2 | 21 |
| 13 | MKUZE | Sep | 1985 | 35.84 | 1.89 | 0.14 | 1.9 | 0.29 | 0.38 | 0.17 | 18 |
| 13 | MKUZE | Dec | 1985 | 40.54 | 1.75 | 0.12 | 1.73 | 0.25 | 0.32 | 0.15 | 17 |
| 13 | MKUZE | Mar | 1986 | 37.1 | 1.15 | 0.1 | 1.1 | 0.2 | 0.32 | 0.14 | 16 |
| 14 | MKUZE | Jul | 1984 | 40.68 | 1.66 | 0.17 | 1.19 | 0.18 | 0.34 | 0.15 | 21 |
| 14 | MKUZE | Dec | 1984 | 37.91 | 1.36 | 0.13 | 1.17 | 0.13 | 0.27 | 0.13 | 15 |
| 14 | MKUZE | Mar | 1985 | 32.78 | 1.72 | 0.11 | 0.96 | 0.23 | 0.35 | 0.14 | 19 |
| 14 | MKUZE | May | 1985 | 30.77 | 1.59 | 0.12 | 0.92 | 0.26 | 0.36 | 0.17 | 23 |
| 14 | MKUZE | Sep | 1985 | 37.6 | 1.71 | 0.13 | 1.6 | 0.22 | 0.32 | 0.16 | 21 |
| 14 | MKUZE | Dec | 1985 | 42.72 | 1.83 | 0.15 | 2.18 | 0.2 | 0.3 | 0.16 | 19 |
| 14 | MKUZE | Mar | 1986 | 34.42 | 1.81 | 0.12 | 1.01 | 0.24 | 0.37 | 0.15 | 20 |
| 15 | MKUZE | Jul | 1984 | 30.82 | 1.55 | 0.12 | 1 | 0.15 | 0.4 | 0.15 | 19 |
| 15 | MKUZE | Dec | 1984 | 37.83 | 1.41 | 0.1 | 0.92 | 0.12 | 0.33 | 0.13 | 14 |
| 15 | MKUZE | Mar | 1985 | 34.51 | 1.44 | 0.11 | 1.16 | 0.22 | 0.28 | 0.12 | 18 |
| 15 | MKUZE | May | 1985 | 30.12 | 1.24 | 0.08 | 0.7 | 0.23 | 0.35 | 0.14 | 16 |
| 15 | MKUZE | Sep | 1985 | 36.02 | 1.43 | 0.1 | 1.29 | 0.22 | 0.32 | 0.14 | 16 |
| 15 | MKUZE | Dec | 1985 | 40.31 | 1.54 | 0.12 | 1.71 | 0.21 | 0.3 | 0.15 | 16 |
| 15 | MKUZE | Mar | 1986 | 41.39 | 1.25 | 0.14 | 1.54 | 0.12 | 0.3 | 0.14 | 18 |
| 16 | MKUZE | Jul | 1984 | 35.58 | 1.86 | 0.12 | 0.94 | 0.25 | 0.47 | 0.18 | 21 |
| 16 | MKUZE | Dec | 1984 | 35.57 | 1.47 | 0.1 | 0.9 | 0.15 | 0.35 | 0.14 | 14 |
| 16 | MKUZE | Mar | 1985 | 31.14 | 1.46 | 0.11 | 1 | 0.23 | 0.3 | 0.14 | 18 |

| | | | | | | | | | | | |
|---|---|---|---|---|---|---|---|---|---|---|---|
| 16 | MKUZE | May | 1985 | 27.21 | 0.94 | 0.08 | 0.55 | 0.25 | 0.28 | 0.17 | 14 |
| 16 | MKUZE | Sep | 1985 | 31.89 | 1.29 | 0.29 | 0.1 | 1.22 | 0.22 | 0.28 | 15 |
| 16 | MKUZE | Dec | 1985 | 35.86 | 1.57 | 0.12 | 1.71 | 0.21 | 0.29 | 0.12 | 17 |
| 16 | MKUZE | Mar | 1986 | 41.49 | 1.54 | 0.13 | 1.73 | 0.17 | 0.34 | 0.13 | 21 |
| 17 | MKUZE | Jul | 1984 | 35.91 | 1.47 | 0.15 | 0.83 | 0.19 | 0.35 | 0.12 | 16 |
| 17 | MKUZE | Dec | 1984 | 37.7 | 1.32 | 0.17 | 1.01 | 0.12 | 0.36 | 0.13 | 18 |
| 17 | MKUZE | Mar | 1985 | 37.45 | 1.88 | 0.18 | 1.35 | 0.24 | 0.38 | 0.12 | 22 |
| 17 | MKUZE | May | 1985 | 28.59 | 1.02 | 0.12 | 0.77 | 0.25 | 0.31 | 0.11 | 21 |
| 17 | MKUZE | Sep | 1985 | 45.17 | 1.99 | 0.2 | 1.97 | 0.23 | 0.44 | 0.14 | 24 |
| 17 | MKUZE | Dec | 1985 | 37.27 | 1.3 | 0.19 | 1.7 | 0.17 | 0.25 | 0.09 | 17 |
| 17 | MKUZE | Mar | 1986 | 43.59 | 1.3 | 0.19 | 1.73 | 0.15 | 0.33 | 0.14 | 21 |
| 18 | HIGHMOOR | Jul | 1984 | 21.58 | 0.5 | 0.03 | 0.33 | 0.08 | 0.3 | 0.08 | 16 |
| 18 | HIGHMOOR | Dec | 1984 | 42.52 | 1.7 | 0.14 | 1.22 | 0.13 | 0.39 | 0.12 | 16 |
| 18 | HIGHMOOR | Mar | 1985 | 27.81 | 1.11 | 0.09 | 0.95 | 0.15 | 0.32 | 0.12 | 12 |
| 18 | HIGHMOOR | May | 1985 | 26.33 | 0.5 | 0.03 | 0.7 | 0.21 | 0.4 | 0.11 | 9 |
| 18 | HIGHMOOR | Sep | 1985 | 24.47 | 0.44 | 0.03 | 0.32 | 0.13 | 0.37 | 0.08 | 13 |
| 18 | HIGHMOOR | Dec | 1985 | 31.48 | 1.28 | 0.1 | 0.93 | 0.18 | 0.3 | 0.12 | 12 |
| 18 | HIGHMOOR | Mar | 1986 | 25.37 | 1.18 | 0.1 | 0.81 | 0.19 | 0.44 | 0.13 | 16 |
| 19 | HIGHMOOR | Jul | 1984 | 21.46 | 0.54 | 0.03 | 0.4 | 0.11 | 0.26 | 0.08 | 19 |
| 19 | HIGHMOOR | Dec | 1984 | 34.02 | 1.52 | 0.12 | 1.12 | 0.11 | 0.34 | 0.11 | 14 |
| 19 | HIGHMOOR | Mar | 1985 | 24.51 | 1.13 | 0.08 | 0.91 | 0.18 | 0.35 | 0.11 | 17 |
| 19 | HIGHMOOR | May | 1985 | 21.26 | 0.74 | 0.06 | 0.85 | 0.18 | 0.28 | 0.1 | 11 |
| 19 | HIGHMOOR | Sep | 1985 | 20.59 | 0.53 | 0.03 | 0.47 | 0.14 | 0.3 | 0.07 | 21 |
| 19 | HIGHMOOR | Dec | 1985 | 28.48 | 0.96 | 0.08 | 1.02 | 0.15 | 0.25 | 0.09 | 11 |
| 19 | HIGHMOOR | Mar | 1986 | 27.26 | 1.02 | 0.08 | 0.84 | 0.16 | 0.28 | 0.09 | 14 |
| 20 | HIGHMOOR | Jul | 1984 | 22.53 | 0.5 | 0.02 | 0.35 | 0.07 | 0.24 | 0.1 | 15 |
| 20 | HIGHMOOR | Dec | 1984 | 32.26 | 1.51 | 0.11 | 1.02 | 0.13 | 0.34 | 0.15 | 12 |
| 20 | HIGHMOOR | Mar | 1985 | 23.9 | 1.22 | 0.09 | 0.81 | 0.19 | 0.37 | 0.13 | 14 |
| 20 | HIGHMOOR | May | 1985 | 21.92 | 0.49 | 0.03 | 0.57 | 0.26 | 0.41 | 0.12 | 14 |
| 20 | HIGHMOOR | Sep | 1985 | 30.6 | 0.86 | 0.05 | 0.77 | 0.22 | 0.29 | 0.1 | 11 |
| 20 | HIGHMOOR | Dec | 1985 | 35.31 | 1.15 | 0.08 | 0.91 | 0.19 | 0.22 | 0.1 | 10 |
| 20 | HIGHMOOR | Mar | 1986 | 27.67 | 0.91 | 0.07 | 0.75 | 0.15 | 0.23 | 0.12 | 10 |
| 21 | HIGHMOOR | Jul | 1984 | 20.07 | 0.45 | 0.02 | 0.19 | 0.07 | 0.34 | 0.09 | 21 |
| 21 | HIGHMOOR | Dec | 1984 | 46.46 | 1.62 | 0.12 | 1.03 | 0.11 | 0.51 | 0.15 | 15 |
| 21 | HIGHMOOR | Mar | 1985 | 27.45 | 1.24 | 0.08 | 0.82 | 0.15 | 0.36 | 0.12 | 13 |
| 21 | HIGHMOOR | May | 1985 | 25.75 | 0.43 | 0.02 | 0.51 | 0.19 | 0.5 | 0.13 | 13 |
| 21 | HIGHMOOR | Sep | 1985 | 24.06 | 0.32 | 0.02 | 0.22 | 0.12 | 0.3 | 0.08 | 10 |
| 21 | HIGHMOOR | Dec | 1985 | 32.29 | 1.39 | 0.1 | 1.09 | 0.17 | 0.29 | 0.13 | 14 |
| 21 | HIGHMOOR | Mar | 1986 | 30.22 | 0.97 | 0.07 | 0.84 | 0.19 | 0.2 | 0.13 | 14 |
| 22 | CATHEDRAL PEAK | Jul | 1984 | 19.76 | 0.37 | 0.01 | 0.15 | 0.07 | 0.26 | 0.09 | 16 |
| 22 | CATHEDRAL PEAK | Dec | 1984 | 34.83 | 1.3 | 0.1 | 0.92 | 0.15 | 0.35 | 0.16 | 15 |
| 22 | CATHEDRAL PEAK | Mar | 1985 | 26.96 | 0.98 | 0.07 | 0.73 | 0.18 | 0.36 | 0.15 | 16 |
| 22 | CATHEDRAL PEAK | May | 1985 | 20.98 | 0.37 | 0.02 | 0.51 | 0.16 | 0.25 | 0.13 | 18 |
| 22 | CATHEDRAL PEAK | Sep | 1985 | 19.71 | 0.33 | 0.02 | 0.26 | 0.1 | 0.23 | 0.08 | 16 |
| 22 | CATHEDRAL PEAK | Dec | 1985 | 23.65 | 0.54 | 0.04 | 0.37 | 0.1 | 0.25 | 0.1 | 14 |
| 22 | CATHEDRAL PEAK | Mar | 1986 | 30.89 | 0.93 | 0.07 | 0.66 | 0.14 | 0.31 | 0.16 | 14 |
| 23 | CATHEDRAL PEAK | Jul | 1984 | 23.46 | 0.44 | 0.01 | 0.16 | 0.08 | 0.23 | 0.08 | 16 |
| 23 | CATHEDRAL PEAK | Dec | 1984 | 35.98 | 1.15 | 0.08 | 0.97 | 0.12 | 0.29 | 0.14 | 12 |
| 23 | CATHEDRAL PEAK | Mar | 1985 | 26.6 | 1.01 | 0.06 | 0.74 | 0.15 | 0.31 | 0.15 | 15 |
| 23 | CATHEDRAL PEAK | May | 1985 | 23.76 | 0.33 | 0.02 | 0.46 | 0.15 | 0.26 | 0.13 | 13 |
| 23 | CATHEDRAL PEAK | Sep | 1985 | 23.79 | 0.39 | 0.03 | 0.36 | 0.09 | 0.21 | 0.1 | 13 |
| 23 | CATHEDRAL PEAK | Dec | 1985 | 33 | 1 | 0.08 | 0.5 | 0.12 | 0.29 | 0.14 | 12 |
| 23 | CATHEDRAL PEAK | Mar | 1986 | 26.88 | 0.9 | 0.05 | 0.7 | 0.15 | 0.3 | 0.18 | 15 |

| | | | | | | | | | | | |
|---|---|---|---|---|---|---|---|---|---|---|---|
| 24 | CATHEDRAL PEAK | Jul | 1984 | 24.53 | 0.42 | 0.01 | 0.12 | 0.07 | 0.19 | 0.08 | 20 |
| 24 | CATHEDRAL PEAK | Dec | 1984 | 38.14 | 1.32 | 0.09 | 0.91 | 0.14 | 0.35 | 0.16 | 14 |
| 24 | CATHEDRAL PEAK | Mar | 1985 | 24.89 | 1.15 | 0.06 | 0.75 | 0.15 | 0.36 | 0.14 | 16 |
| 24 | CATHEDRAL PEAK | May | 1985 | 23.37 | 0.28 | 0.01 | 0.36 | 0.11 | 0.26 | 0.13 | 13 |
| 24 | CATHEDRAL PEAK | Sep | 1985 | 23.67 | 0.35 | 0.02 | 0.24 | 0.09 | 0.23 | 0.09 | 19 |
| 24 | CATHEDRAL PEAK | Dec | 1985 | 25.24 | 0.56 | 0.04 | 0.37 | 0.1 | 0.27 | 0.1 | 18 |
| 24 | CATHEDRAL PEAK | Mar | 1986 | 27.83 | 0.89 | 0.06 | 0.64 | 0.14 | 0.36 | 0.14 | 15 |
| 25 | CATHEDRAL PEAK | Jul | 1984 | 19.13 | 0.41 | 0.02 | 0.22 | 0.06 | 0.22 | 0.07 | 19 |
| 25 | CATHEDRAL PEAK | Dec | 1984 | 36.51 | 1.34 | 0.09 | 1.11 | 0.13 | 0.32 | 0.14 | 14 |
| 25 | CATHEDRAL PEAK | Mar | 1985 | 28.4 | 1.04 | 0.06 | 0.93 | 0.14 | 0.33 | 0.12 | 13 |
| 25 | CATHEDRAL PEAK | May | 1985 | 24.47 | 0.36 | 0.01 | 0.54 | 0.14 | 0.34 | 0.13 | 13 |
| 25 | CATHEDRAL PEAK | Sep | 1985 | 22.32 | 0.35 | 0.02 | 0.22 | 0.11 | 0.27 | 0.07 | 12 |
| 25 | CATHEDRAL PEAK | Dec | 1985 | 24.77 | 0.55 | 0.03 | 0.38 | 0.12 | 0.27 | 0.07 | 13 |
| 25 | CATHEDRAL PEAK | Mar | 1986 | 29.82 | 0.89 | 0.05 | 0.8 | 0.15 | 0.32 | 0.11 | 14 |
| 26 | CATHEDRAL PEAK | Jul | 1984 | 21.46 | 0.44 | 0.01 | 0.21 | 0.07 | 0.2 | 0.1 | 45 |
| 26 | CATHEDRAL PEAK | Dec | 1984 | 35.82 | 1.19 | 0.08 | 0.9 | 0.11 | 0.3 | 0.2 | 13 |
| 26 | CATHEDRAL PEAK | Mar | 1985 | 25.77 | 1.02 | 0.06 | 0.85 | 0.13 | 0.26 | 0.14 | 13 |
| 26 | CATHEDRAL PEAK | May | 1985 | 26.16 | 0.35 | 0.02 | 0.62 | 0.13 | 0.31 | 0.15 | 13 |
| 26 | CATHEDRAL PEAK | Sep | 1985 | 22.74 | 0.42 | 0.01 | 0.26 | 0.09 | 0.24 | 0.1 | 12 |
| 26 | CATHEDRAL PEAK | Dec | 1985 | 24.93 | 0.64 | 0.02 | 0.36 | 0.1 | 0.26 | 0.12 | 13 |
| 26 | CATHEDRAL PEAK | Mar | 1986 | 30.18 | 1 | 0.05 | 0.72 | 0.16 | 0.33 | 0.19 | 16 |
| 27 | CATHEDRAL PEAK | Jul | 1984 | 21.16 | 0.61 | 0.02 | 0.14 | 0.08 | 0.29 | 0.12 | 32 |
| 27 | CATHEDRAL PEAK | Dec | 1984 | 36.88 | 1.35 | 0.08 | 0.85 | 0.13 | 0.32 | 0.22 | 13 |
| 27 | CATHEDRAL PEAK | Mar | 1985 | 28.65 | 1.28 | 0.08 | 0.82 | 0.14 | 0.28 | 0.17 | 13 |
| 27 | CATHEDRAL PEAK | May | 1985 | 25.63 | 0.33 | 0.01 | 0.5 | 0.18 | 0.27 | 0.17 | 12 |
| 27 | CATHEDRAL PEAK | Sep | 1985 | 22.68 | 0.38 | 0.03 | 0.31 | 0.1 | 0.21 | 0.11 | 11 |
| 27 | CATHEDRAL PEAK | Dec | 1985 | 38.57 | 1.45 | 0.1 | 0.94 | 0.15 | 0.29 | 0.2 | 13 |
| 27 | CATHEDRAL PEAK | Mar | 1986 | 29.22 | 0.94 | 0.06 | 0.65 | 0.17 | 0.35 | 0.2 | 13 |
| 28 | UKULINGA | Jul | 1984 | 35.41 | 0.92 | 0.07 | 0.85 | 0.16 | 0.4 | 0.18 | 24 |
| 28 | UKULINGA | Dec | 1984 | 36.5 | 1 | 0.09 | 0.83 | 0.17 | 0.4 | 0.21 | 18 |
| 28 | UKULINGA | Mar | 1985 | 38.2 | 1.77 | 0.12 | 1.15 | 0.15 | 0.5 | 0.24 | 28 |
| 28 | UKULINGA | May | 1985 | 36.03 | 0.68 | 0.05 | 0.83 | 0.16 | 0.43 | 0.19 | 19 |
| 28 | UKULINGA | Sep | 1985 | 34.8 | 1.17 | 0.1 | 0.88 | 0.16 | 0.37 | 0.18 | 30 |
| 28 | UKULINGA | Dec | 1985 | 38.5 | 1.23 | 0.09 | 0.83 | 0.17 | 0.4 | 0.21 | 18 |
| 28 | UKULINGA | Mar | 1986 | 31.37 | 1.13 | 0.09 | 0.97 | 0.18 | 0.28 | 0.19 | 22 |
| 29 | UKULINGA | Jul | 1984 | 37.61 | 1.33 | 0.1 | 0.92 | 0.19 | 0.42 | 0.2 | 12 |
| 29 | UKULINGA | Dec | 1984 | 39.53 | 1.23 | 0.06 | 0.57 | 0.12 | 0.47 | 0.21 | 15 |
| 29 | UKULINGA | Mar | 1985 | 35.81 | 1.43 | 0.08 | 0.7 | 0.17 | 0.5 | 0.23 | 18 |
| 29 | UKULINGA | May | 1985 | 30.84 | 0.7 | 0.04 | 0.68 | 0.19 | 0.42 | 0.2 | 15 |
| 29 | UKULINGA | Sep | 1985 | 44.38 | 1.96 | 0.16 | 1.16 | 0.19 | 0.42 | 0.2 | 23 |
| 29 | UKULINGA | Dec | 1985 | 37.43 | 1.02 | 0.06 | 0.6 | 0.18 | 0.48 | 0.19 | 13 |
| 29 | UKULINGA | Mar | 1986 | 38.76 | 1.43 | 0.09 | 0.86 | 0.19 | 0.37 | 0.21 | 20 |
| 30 | WORLDS VIEW | Jul | 1984 | 30.86 | 1.58 | 0.11 | 0.98 | 0.25 | 0.31 | 0.18 | 18 |
| 30 | WORLDS VIEW | Dec | 1984 | 32.24 | 1.07 | 0.08 | 0.9 | 0.11 | 0.28 | 0.13 | 15 |
| 30 | WORLDS VIEW | Mar | 1985 | 31.28 | 1.41 | 0.1 | 0.8 | 0.16 | 0.29 | 0.14 | 19 |
| 30 | WORLDS VIEW | May | 1985 | 30.51 | 0.86 | 0.04 | 0.61 | 0.25 | 0.32 | 0.18 | 15 |
| 30 | WORLDS VIEW | Sep | 1985 | 47.28 | 2.29 | 0.19 | 1.36 | 0.25 | 0.31 | 0.19 | 21 |
| 30 | WORLDS VIEW | Dec | 1985 | 35.75 | 1.11 | 0.07 | 0.98 | 0.16 | 0.25 | 0.13 | 16 |
| 30 | WORLDS VIEW | Mar | 1986 | 34.28 | 1.26 | 0.08 | 0.82 | 0.15 | 0.29 | 0.16 | 27 |
| 31 | KRANZKLOOF | Jul | 1984 | 37 | 1.6 | 0.11 | 1.16 | 0.19 | 0.31 | 0.18 | 19 |
| 31 | KRANZKLOOF | Dec | 1984 | 31.68 | 1.15 | 0.07 | 0.91 | 0.1 | 0.28 | 0.14 | 17 |
| 31 | KRANZKLOOF | Mar | 1985 | 29.56 | 1.69 | 0.12 | 1.02 | 0.19 | 0.28 | 0.15 | 25 |
| 31 | KRANZKLOOF | May | 1985 | 28.39 | 0.97 | 0.04 | 0.77 | 0.2 | 0.27 | 0.15 | 19 |

| | | | | | | | | | | | |
|---|---|---|---|---|---|---|---|---|---|---|---|
| 31 | KRANZKLOOF | Sep | 1985 | 47.5 | 2.34 | 0.18 | 1.44 | 0.18 | 0.35 | 0.2 | 20 |
| 31 | KRANZKLOOF | Dec | 1985 | 36.54 | 1.18 | 0.07 | 0.96 | 0.15 | 0.25 | 0.15 | 12 |
| 31 | KRANZKLOOF | Mar | 1986 | 33.85 | 1.15 | 0.07 | 0.84 | 0.15 | 0.3 | 0.17 | 27 |
| 32 | KRANZKLOOF | Jul | 1984 | 37.13 | 1.07 | 0.08 | 0.96 | 0.22 | 0.39 | 0.15 | 17 |
| 32 | KRANZKLOOF | Dec | 1984 | 32.79 | 0.96 | 0.06 | 0.91 | 0.09 | 0.32 | 0.12 | 14 |
| 32 | KRANZKLOOF | Mar | 1985 | 34.05 | 1.25 | 0.08 | 0.86 | 0.16 | 0.33 | 0.12 | 20 |
| 32 | KRANZKLOOF | May | 1985 | 34.48 | 0.79 | 0.05 | 0.82 | 0.19 | 0.48 | 0.15 | 17 |
| 32 | KRANZKLOOF | Sep | 1985 | 39.78 | 1.36 | 0.12 | 1.1 | 0.25 | 0.3 | 0.16 | 18 |
| 32 | KRANZKLOOF | Dec | 1985 | 36.91 | 1.2 | 0.09 | 0.95 | 0.19 | 0.34 | 0.1 | 18 |
| 32 | KRANZKLOOF | Mar | 1986 | 37.77 | 1.18 | 0.08 | 0.77 | 0.2 | 0.31 | 0.12 | 29 |

**Supplementary Table 2:** Topsoil (0-200 mm) elements and physical characteristics measured at 31 sites in KwaZulu-Natal (data from Zacharias 1990)

| Site# | K | Ca | Mg | Na | Exchangeable acidity | P | K | Ca | Mg | Na | Exchangeable acidity | Effective cation exchange capacity (meq/100g) | pH | Acid saturation | Field moisture capacity | Sand | Silt | Clay | Organic matter |
|---|---|---|---|---|---|---|---|---|---|---|---|---|---|---|---|---|---|---|---|
| | (meq/100g) | | | | | (mg/kg) | | | | | | | | (%) | ($kg_{water}/kg_{soil}$) | (%) | (%) | (%) | (%) |
| 2 | 0.34 | 13.26 | 8.09 | 0.27 | 0.02 | 0 | 134 | 2658 | 984 | 61 | 2 | 219.8 | 5.3 | 0.1 | 0.4399 | 37.6 | 12.7 | 49.7 | 3.047 |
| 3 | 0.63 | 16.75 | 9.2 | 0.22 | 0.03 | 3 | 247 | 3358 | 1118 | 50 | 3 | 298.3 | 5.3 | 0.1 | 0.4446 | 37 | 8.7 | 54.3 | 3.591 |
| 4 | 1.15 | 9.79 | 3.84 | 0.14 | 0.03 | 0 | 450 | 1963 | 467 | 32 | 3 | 149.5 | 5.1 | 0.2 | 0.4872 | 50.8 | 7.7 | 41.3 | 3.537 |
| 5 | 0.88 | 14.04 | 8.26 | 0.3 | 0.03 | 4 | 343 | 2814 | 1004 | 68 | 3 | 235.1 | 5.3 | 0.1 | 0.4913 | 60.6 | 10.7 | 46.5 | 3.863 |
| 6 | 1.3 | 12.52 | 8.33 | 0.19 | 0.02 | 1 | 508 | 2509 | 1013 | 43 | 2 | 223.6 | 5 | 0.1 | 0.5466 | 33 | 18.2 | 48.8 | 4.97 |
| 7 | 0.42 | 18.32 | 7.53 | 0.28 | 0.04 | 11 | 165 | 3671 | 916 | 65 | 4 | 265.9 | 5.4 | 0.2 | 0.5194 | 32.9 | 23 | 44.1 | 5.205 |
| 8 | 0.98 | 14.53 | 7.92 | 0.14 | 0.03 | 4 | 382 | 2911 | 963 | 32 | 3 | 236 | 5.5 | 0.1 | 0.4708 | 51.6 | 10.8 | 37.6 | 3.41 |
| 9 | 0.42 | 12 | 6.51 | 0.17 | 0.04 | 1 | 165 | 2405 | 792 | 38 | 4 | 191.4 | 5.3 | 0.2 | 0.4851 | 45.8 | 8.4 | 45.8 | 3.718 |
| 10 | 0.31 | 18.42 | 9.16 | 0.25 | 0.03 | 4 | 120 | 3692 | 1113 | 58 | 3 | 271.7 | 5.7 | 0.1 | 0.5203 | 40.2 | 5.8 | 54 | 4.788 |
| 11 | 0.57 | 5.3 | 4.68 | 0.21 | 0.03 | 0 | 224 | 1062 | 569 | 49 | 3 | 107.9 | 5 | 0.3 | 0.3663 | 62.8 | 15.6 | 21.6 | 2.758 |
| 12 | 0.59 | 3.72 | 3.9 | 0.23 | 0.02 | 2 | 231 | 746 | 474 | 52 | 2 | 84.6 | 4.8 | 0.3 | 0.4577 | 55.5 | 22 | 22.5 | 2.485 |
| 13 | 1.58 | 11.65 | 5.55 | 0.07 | 0.02 | 2 | 619 | 2335 | 675 | 15 | 1 | 188.7 | 5.1 | 0.1 | 0.5622 | 39.2 | 17.1 | 43.7 | 4.879 |
| 14 | 1.06 | 6.9 | 5.19 | 0.74 | 0.02 | 1 | 413 | 1383 | 632 | 170 | 2 | 139.1 | 5.2 | 0.2 | 0.4174 | 30.7 | 12.8 | 56.5 | 2.72 |
| 15 | 1.02 | 4.45 | 3.08 | 0.1 | 0.02 | 0 | 398 | 891 | 375 | 22 | 1 | 86.7 | 5.3 | 0.2 | 0.3634 | 71.9 | 1.9 | 26.2 | 1.469 |
| 16 | 0.81 | 3.82 | 2.52 | 0.06 | 0.02 | 0 | 316 | 766 | 307 | 14 | 1 | 72.3 | 5 | 0.2 | 0.3401 | 70 | 2.7 | 27.3 | 2.031 |
| 17 | 1.38 | 10.16 | 5.78 | 0.25 | 0.02 | 3 | 538 | 2036 | 703 | 58 | 2 | 175.9 | 5.3 | 0.1 | 0.477 | 58.6 | 10.1 | 31.3 | 3.083 |
| 18 | 0.23 | 0.77 | 0.35 | 0.02 | 1.44 | 3 | 89 | 154 | 42 | 4 | 130 | 28.1 | 4.3 | 51.5 | 0.6429 | 57.2 | 9.7 | 33.1 | 6.257 |
| 19 | 0.42 | 1.21 | 0.61 | 0.03 | 3.98 | 2 | 164 | 243 | 74 | 7 | 359 | 62.5 | 4 | 63.7 | 0.7767 | 49.2 | 13.4 | 37.4 | 5.985 |
| 20 | 0.22 | 0.87 | 0.39 | 0.02 | 1.92 | 6 | 88 | 174 | 48 | 6 | 173 | 34.2 | 4.2 | 55.9 | 0.7639 | 58.6 | 3 | 38.4 | 7.744 |
| 21 | 0.31 | 0.77 | 0.54 | 0.08 | 0.94 | 1 | 120 | 153 | 65 | 18 | 85 | 26.4 | 4.4 | 35.7 | 0.8664 | 49 | 18.4 | 32.6 | 7.944 |
| 22 | 0.3 | 1.09 | 0.98 | 0.05 | 0.83 | 0 | 118 | 219 | 120 | 11 | 74 | 32.5 | 4.3 | 25.4 | 0.663 | 53.4 | 14.5 | 32.1 | 7.418 |
| 23 | 0.34 | 1.19 | 1.24 | 0.06 | 2.15 | 0 | 133 | 239 | 151 | 13 | 193 | 49.8 | 4.1 | 43.1 | 0.6454 | 17.8 | 19 | 63.2 | 6.91 |
| 24 | 0.28 | 1.26 | 0.93 | 0.04 | 1.02 | 0 | 111 | 253 | 113 | 10 | 92 | 35.3 | 4.3 | 28.7 | 0.665 | 9.4 | 26.9 | 63.7 | 6.638 |
| 25 | 0.24 | 1.51 | 1.49 | 0.03 | 1.6 | 0 | 93 | 303 | 181 | 7 | 144 | 48.7 | 4.3 | 32.9 | 0.8641 | 46.5 | 21.7 | 31.8 | 7.799 |
| 26 | 0.28 | 2.75 | 4.38 | 0.03 | 2.07 | 0 | 110 | 551 | 532 | 7 | 186 | 95.1 | 4.2 | 21.8 | 0.6134 | 30.7 | 14.9 | 54.4 | 5.151 |
| 27 | 0.19 | 1.18 | 1.33 | 0.05 | 1.4 | 0 | 75 | 236 | 161 | 12 | 126 | 41.5 | 4.3 | 33.7 | 0.5614 | 20.6 | 21.2 | 58.2 | 7.127 |
| 28 | 0.19 | 13.44 | 8 | 0.24 | 0.09 | 10 | 74 | 2694 | 973 | 56 | 8 | 475.2 | 4.9 | 0.4 | 0.262 | 15.1 | 20.1 | 64.8 | 4.79 |
| 29 | 0.15 | 8.23 | 5.48 | 0.3 | 0.15 | 1 | 59 | 1649 | 667 | 69 | 14 | 143.1 | 4.6 | 1 | 0.3401 | 34.2 | 28.4 | 37.4 | 4.22 |
| 30 | 0.32 | 1.21 | 1.39 | 0.08 | 1.3 | 1 | 127 | 243 | 169 | 19 | 117 | 43 | 4.3 | 30.2 | 0.4577 | 48.4 | 24.3 | 27.3 | 4.05 |
| 31 | 0.14 | 0.21 | 0.59 | 0.1 | 1.8 | 1 | 55 | 42 | 72 | 24 | 162 | 28.4 | 4.1 | 63.3 | 0.5622 | 14.4 | 7 | 78.6 | 3.64 |
| 32 | 0.11 | 0.29 | 0.35 | 0.03 | 1.54 | 3 | 45 | 58 | 43 | 7 | 138 | 23.2 | 4.1 | 66.2 | 0.4174 | 41 | 49.1 | 9.9 | 1.94 |
